# Immediate effects of light on circadian eclosion and locomotor activity depend on distinct sensory input pathways

**DOI:** 10.1101/2023.04.21.537872

**Authors:** Daniel Bidell, Natalie-Danielle Feige, Tilman Triphan, Claudia Müller, Dennis Pauls, Charlotte Helfrich-Förster, Mareike Selcho

## Abstract

Animals need to be able to sharpen circadian behavioural output in the adaptation to the variable environment. Light is the main entraining signal of the circadian clock, but can also directly increase alertness, locomotor activity, body temperature and heart rate in diurnal animals including humans. Thus, immediate effects of light can enhance or even overwrite circadian output and thereby mask circadian behaviour.

In *Drosophila melanogaster*, immediate light effects are most evident as a lights-on response in two well described behavioural rhythms of the fly – the emergence rhythm of the adult insect from the pupa, called eclosion, and the diurnal rhythm of locomotor activity. Here, we show that the immediate effect of light on rhythmic eclosion depends on the R8 photoreceptor cells of the compound eyes, while the light response of locomotor activity is triggered by different light detecting cells and organs, that seem to compensate for the loss of each other.

## INTRODUCTION

An appropriate daily timing of behaviour is of critical importance for animal fitness ^1,2^. Light shapes the daily timing in two ways: (1) the cyclic change in light intensity acts as an entraining signal, synchronizing the endogenous (circadian) timing system to appropriately adjust an organism to the 24h environmental period; (2) light directly modulates behaviour i.e. increases alertness, locomotor activity, body temperature and heart rate in diurnal animals including humans ^3–7^ and supresses activity and promotes sleep in nocturnal animals ^8,9^. These immediate light effects are independent of the circadian clock and since they often obscure (“mask”) circadian behaviours, they are also referred to as “masking” ^10^. The immediate light effects are essential for appropriate responses of the animal to changes in the environment and for sharpening the behavioural output. Interestingly, light responses are not only shown to be independent of the circadian clock, they can even provoke quasi-wildtype activity rhythms in clock-less and thus arrhythmic fruit flies and mice under natural-like conditions ^11–13^. This clearly underlines their importance.

In *Drosophila melanogaster*, the immediate light effects are most evident as a lights-on response in the two well described behavioural rhythms of the fly – the emergence rhythm of the adult insect from the pupa, called eclosion, and the adult activity rhythm ^14–17^. Eclosion is gated by the circadian clock to the early day, most probably to prevent desiccation and enhance survival rate. A light stimulus induces a rapid increase in eclosion rate (lights-on effect) that is eliminated in flies without eyes and potentially in mutants lacking ocelli ^14,15^. Even though the lights-on effect is gone, eclosion remains synchronized to the light-dark cycle in eyeless flies by the circadian blue-light photopigment Cryptochrome (CRY; ^14,18,19^, reviewed by ^20^). Similarly, the locomotor activity rhythm of adult eyeless flies remains synchronized to the light-dark cycle due to entrainment by CRY (reviewed by ^21^), but the immediate increase of activity after lights-on, also known as “startle response” disappears after elimination of the eyes ^16,22^. Subsequent studies showed that several rhodopsins contribute to this immediate light effect in adult flies ^23,24^.

Flies receive light information via two external organs, the compound eyes and ocelli, as well as the internal extraretinal Hofbauer-Buchner (H-B) eyelets. In addition, a small subset of central brain neurons express the blue-light sensitive photopigment CRY ^18,19^. The fruit fly’s compound eye consists of around 800 ommatidia, each of them equipped with six outer (R1-6) and two inner (R7, R8) photoreceptor cells. While the outer photoreceptor cells express the photoreceptor protein Rhodopsin 1 (Rh1), the inner ones express specific combinations of Rh3, Rh4, Rh5 and Rh6 (reviewed by ^25^). Photoreceptor proteins are specifically expressed in two main ommatidia types: pale ommatidia express Rh3 in R7 and Rh5 in R8 photoreceptors and yellow ommatidia Rh4 in R7 and Rh6 in R8 cells. A third type, the DRA ommatidia, positioned in the dorsal rim area (DRA) expresses Rh3 in R7 and R8 cells. The photoreceptors R1-6 were shown to be involved in dim light vision and the perception of motion ^26^, while R7/R8 are involved in colour vision ^26–28^ and DRA photoreceptors are involved in polarization vision ^29^. Besides the large compound eyes, fruit flies contain three simple dorsal eyes called ocelli. Ocelli consist of only one photoreceptor type, containing Rh2 ^30,31^. The ocelli do not form an image or perceive objects in the environment, instead they are sensitive to changes in light intensities and seem to response to polarized light. As Rh2 is highly sensitive to UV light, the ocelli provide information to distinguishing between sky and the ground (reviewed by ^32^). This enables the fly to maintain its orientation in space. The H-B eyelets evolve from the larval Bolwig’s organ and express Rh6 ^20,33–35^. These extraretinal light detecting cells are involved in entrainment at high light intensities ^36^. In addition to the light detecting organs, *Drosophila* contains CRY expressing neurons in the brain and compound eyes, sensitive to blue light ^37–40^. Beside the compound eyes, CRY is the main photoreceptor involved in light entrainment of the circadian clock ^16,41^. In our study, we aim to identify the neuronal correlates of the immediate light effects that trigger specific behavioural responses counteracting or reinforcing circadian rhythms. For this, we use eclosion and locomotor activity as behavioural readouts to examine immediate light effects on circadian behaviours. We provide evidence that the light effect on eclosion is mediated by Rh5-positive R8 photoreceptor neurons of the compound eyes. In contrast, the ocelli, the H-B eyelets, as well as light sensitive CRY-positive cells are not required to elicit this lights-on response. Interestingly, the light response on locomotor activity in the night remains in flies without eyes, ocelli or CRY-positive cells. We hypothesize redundant signalling pathways and propose that light perceiving cells and organs compensate the loss of each other, enabling flies to react to changes in their environment. Thus, immediate effects of light on circadian behaviours seem to depend on different underlying neuronal networks.

## RESULTS

Light elicits an immediate increase in eclosion at lights-on in fly populations monitored under 14:10 light:dark cycles (Fig.1A). This immediate effect is also visible in flies kept under constant darkness when a light pulse is given at times around circadian time 0 (CT0, Fig.1B-D). In detail, in line with previous studies ^15^, we could show that a twenty minutes white light pulse (I = 4,1 W/m^2^) one hour before or after CT0 elicits an immediate eclosion response in control flies (Fig.1B-D). The light response thus enables the flies to eclose promptly in response to their environment, reinforcing the behaviour controlled by the clock. Next, we tested whether the pigmentation of the eyes plays a role in the detection of light necessary to elicit immediate eclosion. For this, we monitored eclosion in red-eyed CantonS (CS) and white-eyed (*w*^*1118*^) flies. Both groups showed a clear eclosion peak in response to light, even though the distribution within the twenty minutes is different (Fig.1C,D, S1). Nevertheless, eye-colour does not influence the immediate response to light. Further, we tested different mutants to discover the light perceiving cells and organs triggering immediate eclosion. The lights-on response is gone in flies lacking the compound eyes (*cli*^*eya*^, Fig.2B; ^14,15^) and in *norpA* (*no receptor potential*) mutants with disturbed phospholipase C function (*norpA*^*p41*^, Fig.2C). In contrast, flies with disabled light perception in photoreceptors of the ocelli (*rh2*^*1*^, Fig.2D), *cry*-positive cells (*cry*^*01*^, Fig.2E) or the Rh6-positive H-B eyelets (*rh6*^*1*^, Fig.S2), respectively, showed an immediate increase in eclosion in response to light indicating the exclusive requirement of the compound eyes in the immediate effect of light on eclosion.

**Fig.1:**
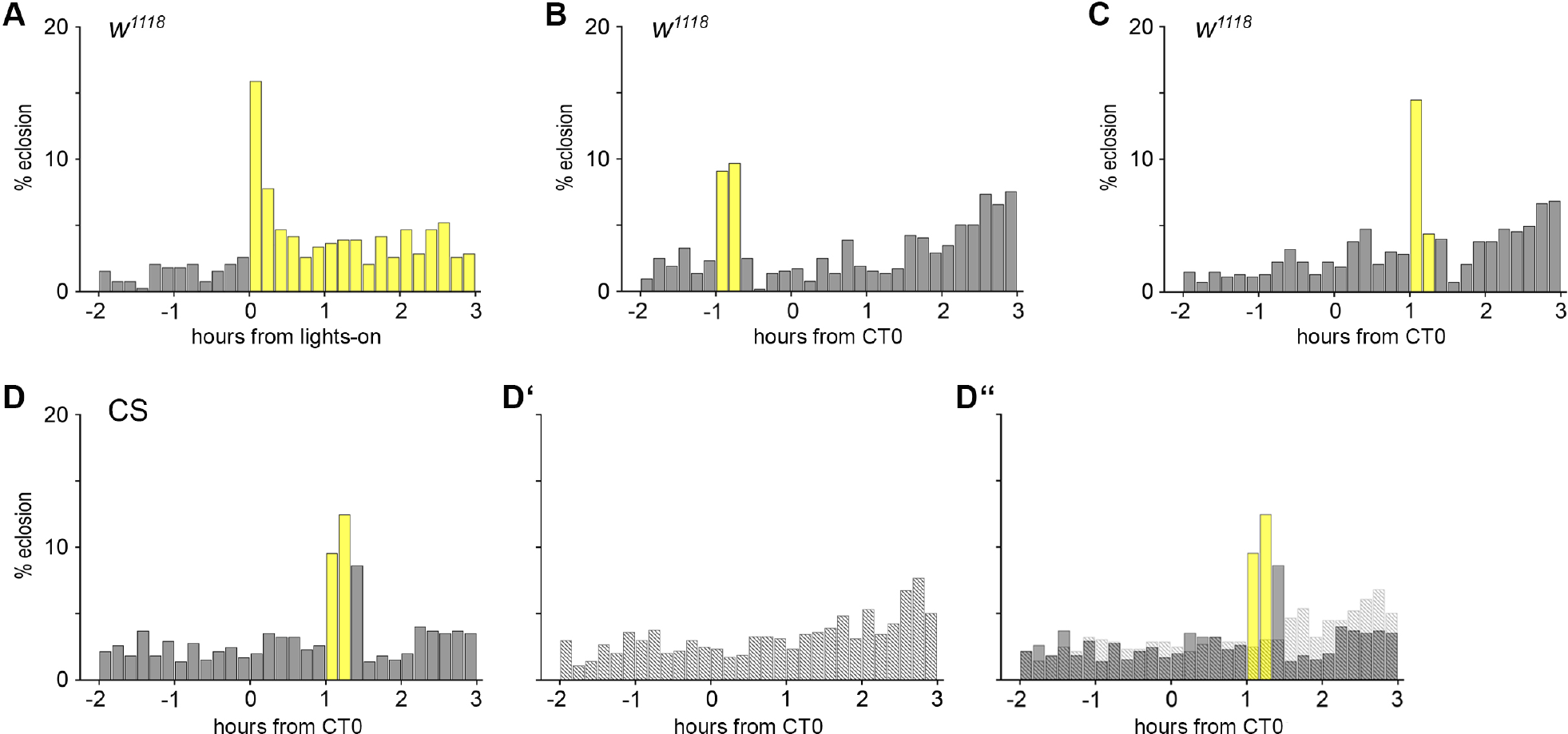
The immediate light effect on eclosion behaviour. **(A)** Eclosion pattern of *Drosophila* flies in ten minutes intervals at the times around lights-on. Light elicits an immediate increase in eclosion. **(B**,**C)** The lights-on response is visible in flies perceiving a twenty minutes light pulse (B) one hour before (-1) or (C) one hour after (1) expected lights-on at CT (circadian time) 0. **(D-D’)** Wildtype CantonS (CS) flies show an immediate response to light (D), while flies in darkness lack the eclosion peak (D’). (D’’) The third plot combines (D) and (D’) to visualize the immediate light response in comparison to the appropriate controls monitored in darkness. Grey bars: % eclosion in dark phase, yellow bars: % eclosion in light phase; dashed bars: % eclosion in controls. n= 384 (A), 516 (B), 524 (C), 649 (D), 636 (D’).

**Fig.2:**
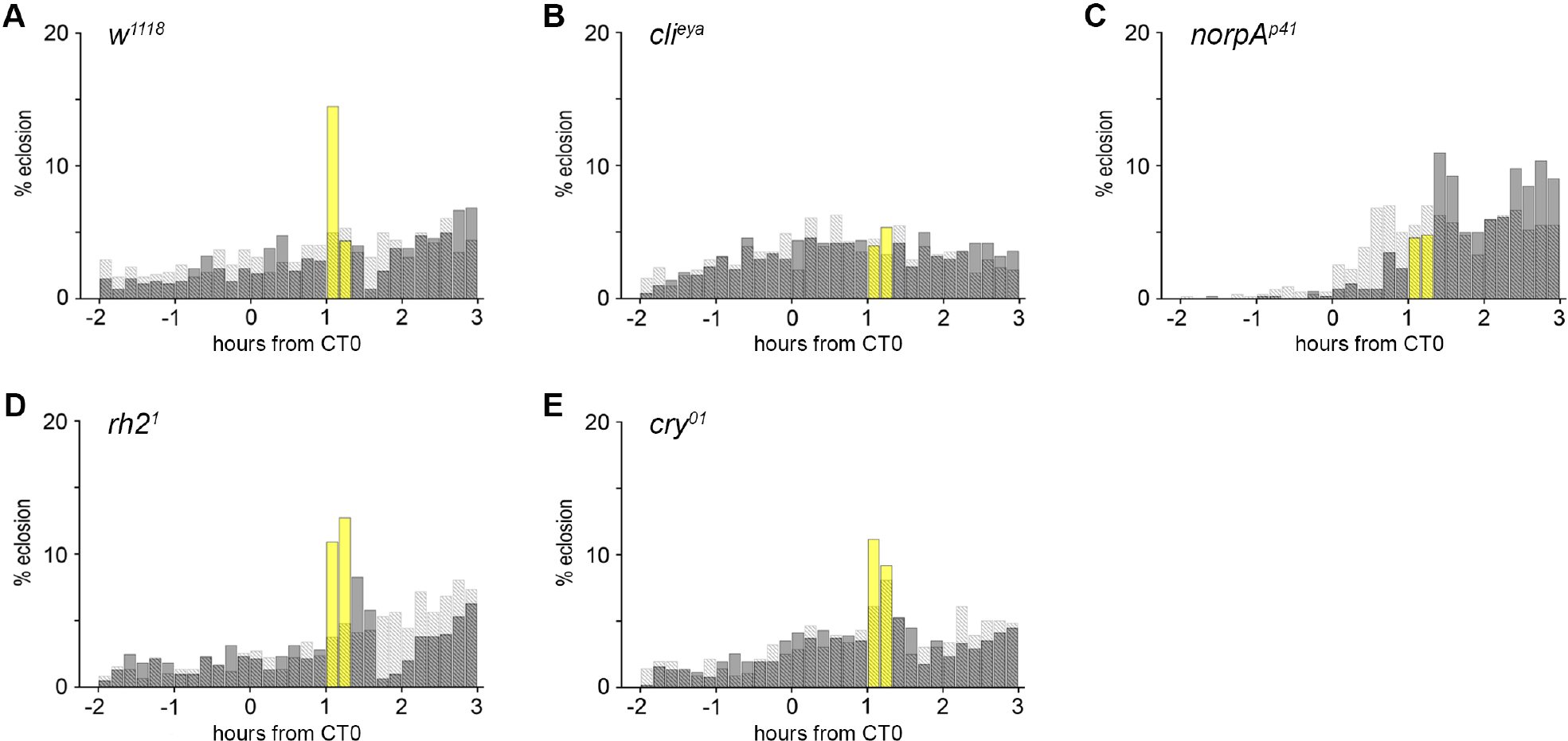
The immediate light effect on eclosion behaviour requires the compound eyes and phospholipase C activity. **(A-E)** Eclosion pattern in ten minutes intervals at the times around circadian time (CT) 0. Each plot visualizes the results for the experimental (grey, yellow bars) and control groups (dashed bars) as shown in Fig.1D-D’’. **(B**,**C)** Flies without eyes (B, *cli*^*eya*^) and impaired phospholipase C activity (C, *norpA*^*p41*^) lack the lights-on response. **(D**,**E)** Flies lacking the rhodopsin (Rh) of the ocelli photoreceptor cells (D, *rh2*^*1*^) or the photoprotein Cryptochrome (E, *cry*^*01*^) respond to light. Grey bars: % eclosion in dark phase, yellow bars: % eclosion in light phase; dashed bars: % eclosion in darkness controls. n_exp_, n_ctrl_ = 524, 544 (A); 502, 508 (B); 520, 539 (C); 604, 584 (D); 510, 556 (E).

Our data so far suggest that the compound eyes are required for the immediate light effect on eclosion. Light information is received by photoreceptor cells of the compound eyes and converted into neuronal activity. The outer photoreceptor cells R1-R6 contain Rh1, the inner R7 cells express Rh3 (R7pale) or Rh4 (R7yellow) and proximal R8 cells express either Rh5 (R8pale) or Rh6 (R8yellow), while DRA ommatidia express Rh3 in R7 and R8. Thus, the different photoreceptor cells respond to light of a specific spectrum as the five different rhodopsin’s of the compound eyes have different absorption spectra ^42–44^. Therefore, illumination by monochromatic light of different wavelength will allow to address specific sets of Rh cells only and thereby give insights into the sufficiency of the different photoreceptor cells in this context. Flies showed an immediate behavioural response to twenty minutes blue (455– 475 nm, I = 3,6 W/m^2^), green (510– 545 nm, I = 2 W/m^2^) and red (625– 642 nm, I = 2,3 W/m^2^) light pulses (Fig.3A-C). In contrast to mammals, where immediate light-effects depend exclusively on blue light-sensitive melanopsin-expressing retinal ganglion cells^45–49^, flies are able to additionally respond to longer wavelengths. As red light of 633nm could only be absorbed by Rh1 or Rh6, we monitored the immediate light responses in flies without Rh1 (*ninaE*^*17*^), flies that lack Rh6 (*rh6*^*1*^) and double mutants without both of these rhodopsins (*ninaE*^*17*^; *rh6*^*1*^). As expected, the lights-on response is missing in the *rh1,rh6* double mutant (Fig.3F). Lacking just one of the red light detecting rhodopsins leads to a reduced lights-on response (Fig.3D,E). Interestingly, however, flies lacking Rh1 and Rh6 show an immediate behavioural response to a twenty minutes white light pulse (I = 4,1 W/m^2^, Fig.3G), indicating that Rh1 and Rh6 are both involved in the immediate effect of red light, but are dispensable for the immediate effect of white light on eclosion. In addition, as Rh1 is the only rhodopsin expressed in the outer photoreceptor cells, R1-R6 might not be required for the lights-on response to white light. To further disentangle the role of the R7, R8 cells, we screened rhodopsin and photoreceptor mutants for their behavioural response to white light (Fig.3G-J). The quadruple mutant (*rh5*^*2*^*;rh3*^*1*^,*rh4*^*1*^,*rh6*^*1*^) lacks all rhodopsins of the inner photoreceptors, so that light perception is only possible via R1-R6. The lack of the lights-on response (Fig.3H) indicates that (1) R7 and/ or R8 transmit light information for the lights-on response and (2) functional outer photoreceptors are not sufficient to trigger immediate eclosion. To address if R7 is involved in the immediate effect of light, we monitored eclosion in *sevenless* mutants (*sev*^*LY3*^, Fig.3I). Flies without R7 photoreceptor cells show an immediate response, so that R8 cells are considered for the transmission of light information. R8 cells express Rh5 or Rh6. Since the absence of Rh6 has no effect on the immediate response to light (Fig.3G and Fig.S2), we tuned our attention to the role of *rh5* mutants on eclosion (*rh5*^*2*^, Fig.3J). As expected, flies lacking Rh5 show no lights-on response. Thus, pale R8 ommatidia expressing functional Rh5 turned out to be essential for the immediate response to light. In line with this hypothesis, we optogenetically activated *rh5*-expressing R8 neurons (Fig.3K,L). Photostimulation of *rh5*-positive neurons via Channelrhodopsin-2^XXL^ (*rh5*^*G*^*>chop2*^*XXL*^, Fig.3K; ^50,51^) using a blue light pulse triggered immediate eclosion, while the response is absent in control flies (Fig.3L). This demonstrates that activation of *rh5*-positive R8 cells triggers immediate eclosion. Our data demonstrate that Rh5-expressing R8 cells are necessary and sufficient for the immediate light effect on eclosion.

**Fig.3:**
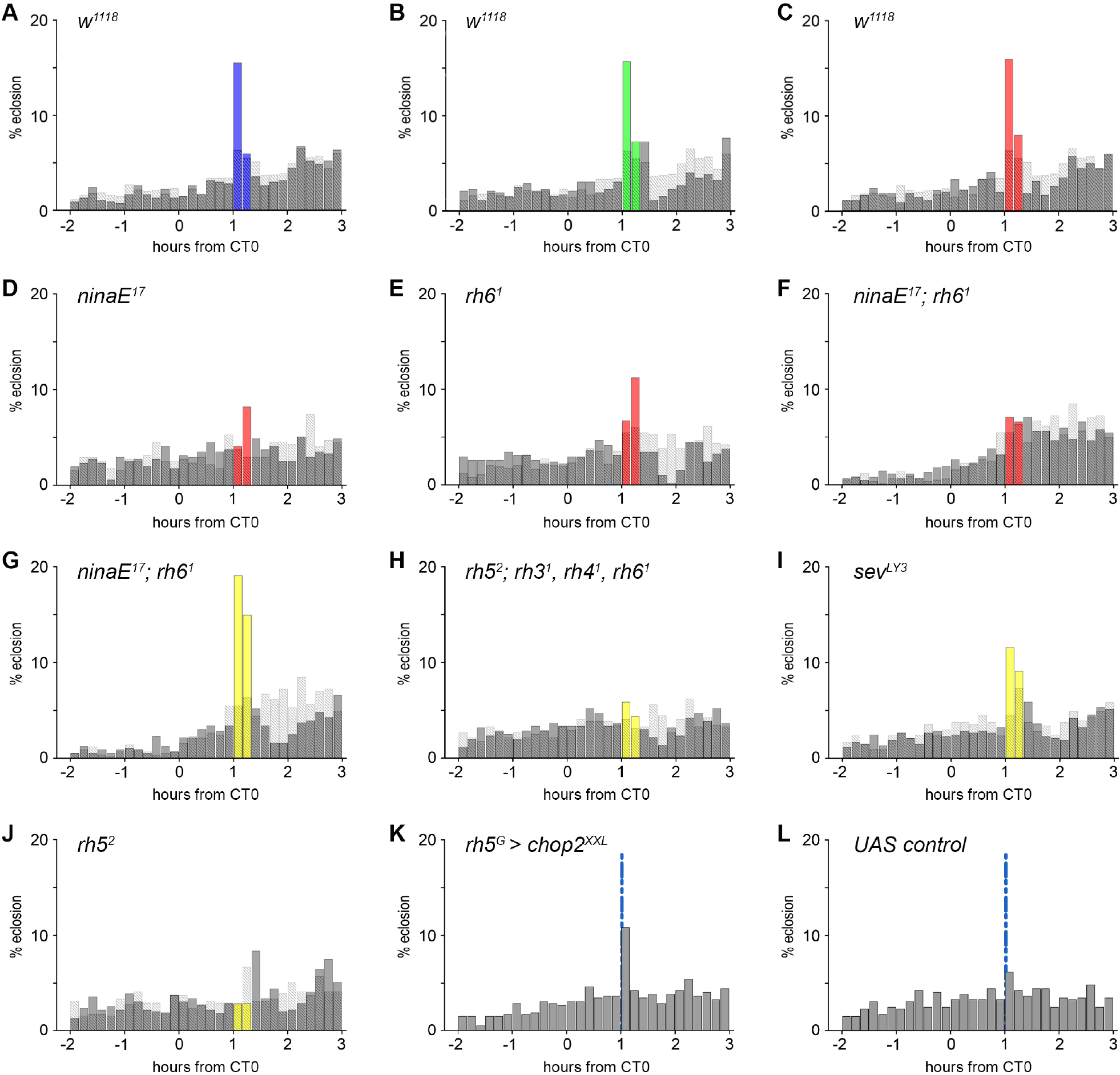
The immediate light effect on eclosion behaviour depends on R8 cells. **(A-J)** Eclosion pattern in ten minutes intervals at the times around circadian time (CT) 0. Each plot visualizes the results for the experimental (grey, yellow bars) and control groups (dashed bars) as shown in Fig.1D-D’’. **(A-C)** Twenty minutes blue (A, 455– 475 nm, I = 3,6 W/m^2^), green (B, 510– 545 nm, I = 2 W/m^2^) and red (C, 625– 642 nm, I = 2,3 W/m^2^) light pulses elicit immediate eclosion. **(D-F)** The eclosion response to red light is visible in *rh1* (D, *ninaE*^*17*^) and *rh6* mutants (E, *rh6*^*1*^), but gone in *rh1,rh6* double mutants (F, *ninaE*^*1*^; *rh6*^*1*^). **(G)** In contrast, *rh1,rh6* double mutants (*ninaE*^*1*^; *rh6*^*1*^) respond with an increase of eclosion to white light (I = 4,1 W/m^2^). **(H)** The quadruple mutant (*rh5*^*2*^; *rh3*^*1*^, *rh4*^*1*^, *rh6*^*1*^), lacking all rhodopsins of the inner photoreceptors, shows no reaction to light. **(I)** The lights-on response in flies without R7 cells (*sev*^*LY3*^). **(J)** Flies lacking Rh5 (*rh5*^*2*^) do not respond to light with increased eclosion. **(K)** Optogenetic activation of *rh5*-positive neurons with a 2 min blue light pulse (blue line; 455-475nm, I = 3,41 μW/mm^2^) one hour after expected lights-on elicits eclosion. **(L)** Control flies without Gal4 expression (*w*^*1118*^; *chop2*^*XXL*^) do not respond to the 2 min light pulse. n_exp_, n_ctrl_ = 534, 1543 (A); 521, 1543 (B); 547, 1543 (C); 511, 512 (D); 579, 731 (E); 604, 567 (F); 561, 567 (G); 595, 518 (H), 525, 531 (I); 585, 506 (J); n= 516 (K), 515 (L).

Eclosion is not the only circadian behaviour that can be modulated by light. To explore whether immediate light effects are generally mediated by R8 cells of the compound eyes as demonstrated above, we turned our attention to locomotor activity. Locomotor activity in *Drosophila* is regulated by the circadian clock so that flies display low levels of nocturnal activity and higher levels of diurnal activity. Importantly, flies display a lights-on response in the transition from night to day at ZT0 (startle response, Fig.4A). To investigate immediate light effects, we applied a twenty minutes white light pulse (∼435 – 780 nm, I = 0.923 W/m^2^) at ZT22, two hours before the anticipated lights-on response in the transition from night to day. Former studies clearly showed that unexpected light at night elicits an immediate increase in activity ^52–54^. As expected, the light pulse elicited an immediate increase in locomotor activity, similar to the light effect on eclosion (Fig.4A, Fig.1). Interestingly, in contrast to eclosion, the immediate light response in locomotor activity is also present in eyeless flies that lack additionally functional CRY (*cli*^*eya*^*;cry*^*b*^; Fig.4B). Thus, neither photoreceptor cells of the compound eyes nor CRY are required for the increase in activity in response to light (Fig.4B and Fig.S3 for *rh* and *norpA* mutants). In addition, we analysed locomotor activity in two different *cry* mutants to clarify the role of photosensitive neurons outside the visual system (*cry*^*b*^ and *cry*^*01*^). Flies without functional CRY responded to light at ZT22 with an immediate increased locomotor activity (Fig.4C and Fig.S3A). Further, immediate light effects can be seen in flies with disabled light perception in photoreceptors of the ocelli (*rh2*^*1*^, Fig.4D) and flies without Rh5 and Rh6 and therefore without functional eyelets (*rh5*^*2*^, *rh6*^*1*^; Fig.4E). Thus, all the different cells and organs that perceive light appear to mediate its immediate effects and compensate for the failure of individual photoreceptors, so that flies can respond immediately to a light stimulus. In contrast, flies without a functional histidine decarboxylase (*hdc*^*JK910*^), the enzyme necessary for histamine synthesis, show no increase in locomotor activity in response to light (Fig.4F). Histamine is the main transmitter of the photoreceptor cells of the eyes, ocelli and eyelets ^55^, leaving the hypothesis that *cry*-positive cells alone may not be sufficient to elicit the lights-on response.

**Fig. 4:**
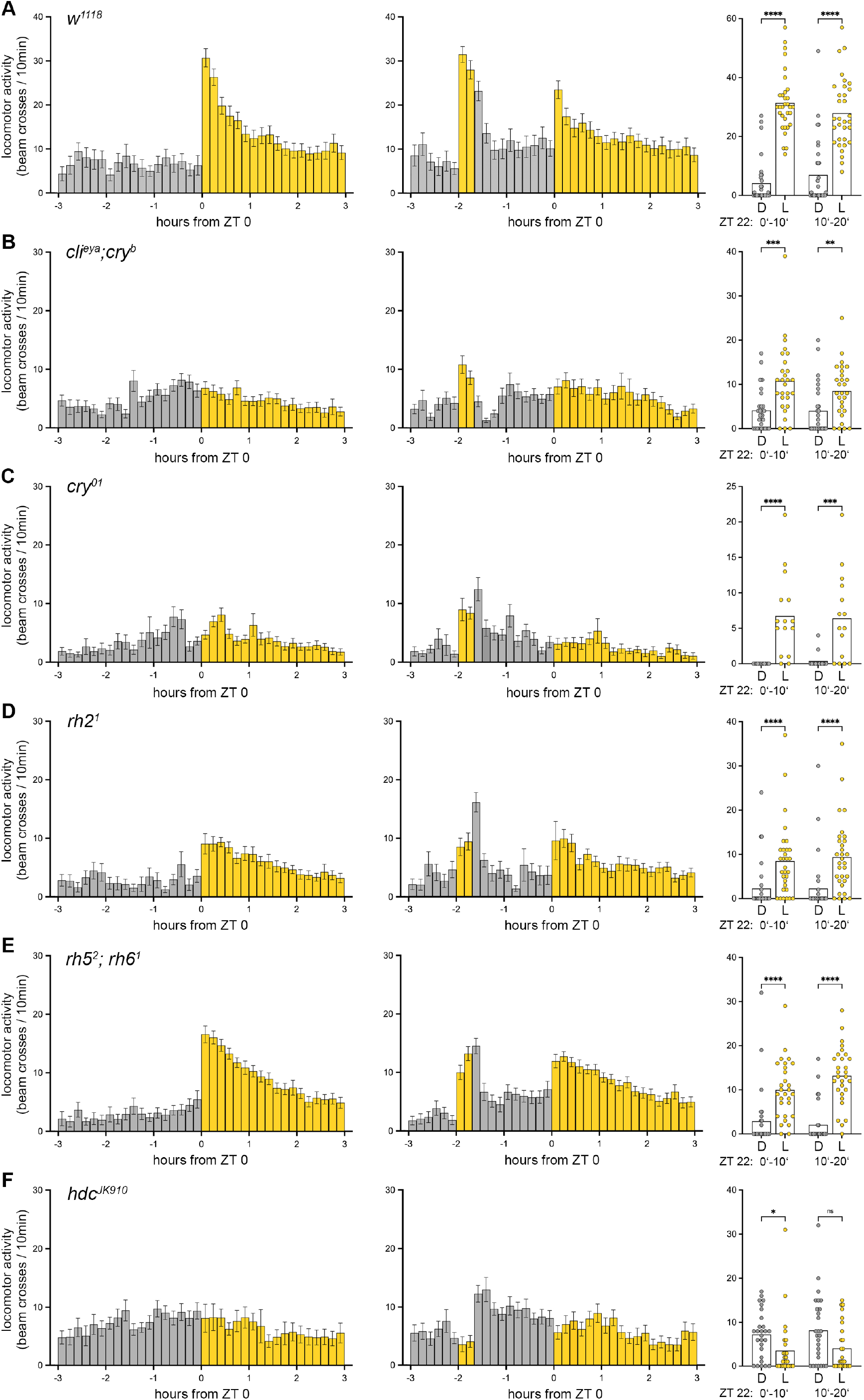
The immediate light effect on locomotor activity is visible in flies without functional eyes, photosensation in *cry*-positive cells or ocelli. **(A-F)** Activity pattern in ten minutes intervals at the time around Zeitgeber time (ZT) 0. First and second column show bar plots of mean ± SEM activity at the day the light pulse was applied (second column) and the activity of the same flies on the previous day (first column). The third column visualizes the comparison between the mean activity at ZT22 in 10 minutes intervals (0’-10’ and 10’-20’) during the light pulse (L) and the previous control day in darkness (D). **(A)** Activity data of control (*w*^*1118*^) and **(B)** eyeless (*cli*^*eya*^; *cry*^*b*^) flies, **(C)** flies lacking Cryptochrome (*cry*^*01*^), **(D)** flies without functional ocelli photoreceptors (*rh2*^*1*^), **(E)** flies without Rh5 and Rh6 (*rh5*^*2*^; *rh6*^*1*^) do respond to light. **(F)** The light response is absent in flies that lack the histidine decarboxylase (*hdc*^*JK910*^) and therefore histamine, the transmitter of photoreceptor cells. n = 27-32; asterisks denote level of significance: ^**^p≤0.01, ^***^p≤0.001, ^****^p≤0.0001.

In summary, our data provides evidence that immediate light effects are present on eclosion and locomotor activity. However, the underlying signalling pathways appear to be different.

## DISCUSSION

Even though light triggers immediate eclosion and activity, modifying circadian behaviour, the necessary underlying networks seem to differ already at sensory input level. The lights-on response on eclosion depends on Rh5-positive R8 cells of the compound eyes, while the light response on locomotor activity works without eyes, ocelli, eyelets or photosensitive *cry*-positive brain cells and is only absent when histamine is missing. For activity increase at night the different light detecting organs and cells can compensate the loss of each other, probably to ensure the ability to react to the unexpected external stimulus.

The functional redundancy of photoreceptors was also described in nocturnal mice. Here, bright light at night leads to an inhibition of locomotor activity, whereas dim light increases activity ^56–58^. Mice with degenerated retina (*rd/rd* mice), devoid of rods and cones, lack the increase of activity by dim light, but still show activity inhibition by bright light. In addition, mice lacking the photopigment melanopsin (*Opn4*^*-/-*^) in the retinal ganglion cells also show inhibition of activity by bright light, while double mutants (*Opn4*^*-/-*^; *rd/rd*) loose the ability to respond to light ^47,48,59^. Thus, also in mice, several photopigments contribute to the immediate light responses and these are able to compensate to some degree the loss of each other to ensure immediate responses to unexpected external stimuli.

The immediate effect of light on locomotor activity is visible every morning in flies under LD rhythms at ZT0 (16,17). Interestingly, former studies showed, that the immediate increase in locomotion (= startle response) at ZT0 depends on the compound eyes and is completely absent in eyeless flies^16^. Here, we observed the same: eyeless flies (*cli*^*eya*^*;cry*^*b*^ mutants, which lack functional CRY in addition to the eyes) as well as flies without histamine (*hdc*^*JK910*^ mutants) lack the startle response at ZT0 (first column, Fig.4B,F). Nevertheless, a light pulse during the night two hours before lights-on at ZT22, provokes an immediate increase in activity in *cli*^*eya*^*;cry*^*b*^ mutants (Fig.4B). Only the disruption of neuronal communication of photoreceptor cells of the eyes, ocelli and eyelets by the loss of histamine (*hdc*^*JK910*^ mutants) prevents a startle response at night (ZT22, Fig. 4F). The immediate increase in activity in response to light at night at ZT22 is thus mediated by functionally redundant photoreceptor cells, whereas the startle response at ZT0 depends purely on the receptors in the eyes.

Interestingly, recent data suggests that CRY suppresses activity during the night in flies, as the absence of CRY enhances activity during moonlit nights ^60^. Flies were as active during moonlit nights as they were during the day, suggesting that CRY is important for distinguishing nocturnal low light of moonlight intensity from day light, and most interestingly this was also true for marine bristle worms ^60^. If CRY is present, it seems to suppress activity during the night in the diurnal *D. melanogaster* and to suppress swarming activity during the day in the nocturnal marine bristle worms. This finding for *Drosophila* is corroborated by the present study: *cry*^*b*^ mutants show strong immediate light effects upon a light pulse, but flies that lack the function of all photoreceptors except of CRY (*hdc*^*JK910*^ mutants) don’t show this response. In summary, we conclude that the immediate light effects of adult flies in response to nocturnal light are mediated by all rhodopsins (those in the compound eyes, the H-B eyelets and putatively also the ocelli), while they are inhibited by CRY. This is different for the startle response at ZT0. Here we could not see any inhibiting effect of CRY.

The immediate effect of light on locomotion in *Drosophila* is modulated in a time-of-day-dependent manner: light at night increases activity, while activity is suppressed during the day ^52,54^. The switch in behaviour seems to depend on the daily morphological changes of central clock neurons ^54^ and on a light-mediated circuit switching in the *Drosophila* neuronal clock network ^61^. The differences in the photoreceptor necessity at ZT22 and ZT0 might reflect the changes in time of the day.

Even though the context dependent behavioural plasticity depends on the circadian clock, the increase in activity in response to light is still visible in flies without functional internal clocks ^52,54,62^. The masking effect is therefore clock independent, but we cannot completely exclude that contextualization of the light stimuli is disturbed in *hdc*^*JK910*^ mutants leading to a decrease in activity in response to light at ZT22. In any case, the experiments using the *hdc*^*JK910*^ mutants show that the immediate effects of light on locomotion are mediated via neurotransmission through histamine, which is the main transmitter of the compound eyes, ocelli and H-B eyelets ^55^. The H-B eyelets use additionally acetylcholine (ACh) as a neurotransmitter ^36^ and a very recent paper shows that this is also true for the inner photoreceptors R8 (Xiao et al. 2023, Nature, in press). Most interestingly, ACh appears to transmit photic input to the circadian clock, while histamine transmits visual signals for image detection and motion. Thus, consistent with our results histamine seems to be the transmitter for the direct light responses.

For the immediate effect of light on eclosion, only Rh5 in R8 cells of the compound eyes is needed. Possibly, not all photoreceptors are yet mature at the time of eclosion. This is certainly true for the H-B eyelets that are fully functional only several days after eclosion ^63^. For the other photoreceptors such a delayed maturation is not known, but it is possible that not all the connections to the central brain, in particular those that connect to activity-promoting centres, are fully established.

## METHODS

### Fly stocks

Flies were raised on standard cornmeal and molasses medium at 25°C and 65% relative humidity at a 14:10 light-dark (L:D) cycle unless otherwise stated. For optogenetic experiments, vials have been covered with a light filter foil (Nr. 026; LEE Filters Worldwide, UK). The following flies were used in this study: *cli*^*eya*^ (^64^), *cry*^*01*^ (^65^), *cry*^*b*^ (^66^, kind gift of R. Stanewsky), *hdc*^*JK910*^ (^67^), *nina*^*E17*^ (^68,69^), *norpA*^*p41*^ (^70,71^, kind gift of R. Stanewsky), *rh2*^*1*^ (^72^, kind gift of C. Montell), *rh5*^*2*^ (^73^, kind gift of R. Stanewsky), *rh6*^*1*^ (^74^, kind gift of C. Montell), *nina*^*E17*^; *rh6*^*1*^ (^75^), *sev*^*LY3*^ (^76^) and the following combinations of the mutants were used: *cli*^*eya*^; *cry*^*b*^ and *rh5*^*2*^; *rh6*^*1*^ and *rh5*^*2*^; *rh3*^*1*^, *rh4*^*1*^, *rh6*^*1*^. The Gal4- and UAS-lines used: *rh5*^*G*^ (^72^, kind gift of C. Montell; BDSC #66671), *UAS-chop2*^*XXL*^ (^51^), *10xUAS-IVS-myr::GFP* (^77^; BDSC #32197).

### Eclosion behaviour

To investigate the immediate light effect on eclosion behaviour, an eclosion monitor based on the WEclMon-System ^78,79^ was used (Fig.S4A). Experiments were performed under constant temperature (24.5°C ± 0.2°C) and humidity (around 65% RH). Temperature fluctuations upon light exposure were below 0.2°C. 7 to 9 days old pupae were placed onto a transparent acrylic plate. This plate was placed on an area light with an RGB or White-LED illumination (LED-color: λ(blue)= 455 – 475 nm, λ(green)= 510 – 545 nm, λ(red)= 625 – 642 nm; white LED (λ= ∼430 – 730 nm, Fig.S4B); Hansen GmbH, Germany, Luminous Panel (RGBW)) and an additional IR LED strip was installed at the bottom of the lighting unit (SOLAROX® LED, λ(infrared) = 850nm, IR1-60-850, Winger Electronics GmbH & Co. KG, Germany) and covered with an aluminum box. Flies were kept in darkness and received a twenty minutes light pulse one hour after expected lights-on (CT1) on the second day, or remained in darkness (darkness control). The eclosion monitor is equipped with a camera (DMK 27AUC02 with a TPL 0420 6MP objective; DMK 37BUX287 with a TPL 0620 6MP objective, The Imaging Source Europe GmbH) that took images every two minutes (IC Capture 64bit, V2.5.1547.4007, The Imaging Source Europe GmbH, Germany). We developed a Python-script for Fiji (V2.9.0, ^80^) that was able to independently scan a selected image sequence for eclosion events (detailed description below).

### Eclosion data analysis

The detection of hatching events is based on the difference in brightness between a pupa and an empty pupal case. The latter is almost transparent while a late pupa is considerably darker (Fig.S4C, empty pupal case marked with an asterisk). In the first step of the analysis all pupae are detected. The pupae are darker than the background and can be easily extracted in a single image. After setting a threshold (Fig.S4D), objects are detected, and their outlines stored. Objects that are outside of the parameters defined for a pupa (area too big or too small) are excluded. Note the empty pupal case that is excluded from further analysis. As the pupae are stationary, we can follow them through time and calculate the median grey value of the area contained in their respective outlines (Fig.S4E) for each frame. Alternatively, the average or mode (i.e. most frequent) grey values can be used. A large jump in brightness indicates an eclosion event (Fig.S4F-H). The median grey values are more than doubled from around 40 to almost 100. For each eclosion event a few frames before and after the event are extracted to help with manual confirmation of real hatching events or to exclude inaccurate detections (Fig.S4G). The time for each eclosion event is stored in a csv file for further analysis. Additional checks are implemented to reduce the number of incorrect detections and handle changes in lighting. See the commented source code for details. The complete workflow is implemented as a Python script for Fiji. Specific parameters that depend on camera resolution, optics and lighting can be set manually (e.g. area of the pupae in pixel, difference in brightness for full and empty pupae, etc.). All code and further information can be found at https://github.com/triphant/EclosionDetector.

For the evaluation of eclosion events, a time window of five hours was chosen, from two hours before expected lights-on to three hours after expected lights-on. To calculate the eclosion percentage (% eclosion), the number of flies eclosed in a 10 min time interval was normalized to the number of flies eclosed during the 5 h time window. The bar plots in Fig.1D’’, Fig.2 and Fig.3 visualize the eclosion of flies perceiving a 20 min light pulse and of appropriate control flies kept in darkness (Fig.1D-D’’). n refers to the number of individuals tested.

To analyze changes in eclosion rate in response to the light the eclosion rate for experimental (L) and control (D) flies in the first 10 min (0’-10’) and second 10 min (10’-20’) interval after lights-on at CT1 was calculated:

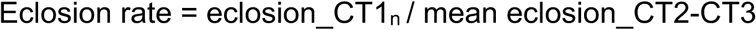

with eclosion_CT1_n_ = eclosed flies at CT1 in the first 10min (0’-10’) or second (10’-20’) ten minutes in light (L) or darkness (D) and eclosion_CT2-CT3 = mean eclosion from CT2 to CT3.

Data was analysed with Excel (Microsoft) and Prism 8.2 (GraphPad). The Shapiro-Wilk test was used to analyse normal distribution and data were compared by an unpaired two-tailed t-test. Not normally distributed data were compared by a nonparametric Mann-Whitney rank sum test. Prism was used to plot the results (Fig.S1,S2). Significance levels refer to the raw p values obtained in the statistical tests.

### Optogenetics

For optogenetic activation, Channelrhodopsin-2^XXL^ (UAS-*chop2*^*XXL*^) has been used to depolarize *rh5*-positive neurons by blue light. Pupae have been collected under red light during their subjective day to monitor eclosion. One hour after expected lights-on pupae received a 2 min blue light stimulus (455 – 475 nm, I = 3,41 μW/mm^2^) and eclosion was monitored and calculated as described above. n refers to the number of individuals tested.

### Locomotor Activity

Flies were entrained to a 12:12 LD rhythm to be able to compare results to the data by ^52,54^. To investigate the lights-on effect on locomotor activity, we placed two to four days old flies in small glass tubes with 2% agarose and 4% sugar on one side into the *Drosophila* Activity Monitoring System (DAM, V2, TriKinetics Inc., USA) and recorded locomotor activity for the next four night-day cycles under constant temperature and humidity (24.9 ± 0.1°C, ∼65% RH). In the first three night-day cycles the light regime did not differ from the entrained rhythm (12:12 LD). On the fourth night, a 20 min light pulse (λ= ∼435 – 780 nm, I = 0.923 W/m^2^) was given two hours before normal lights-on at ZT22. The data received from the DAM system was analyzed by taking the number of counts of a 10 min interval of each tube and calculating the average activity. This was done for both the 6 h time window (-3 h to +3 h from ZT0) in which the 20 min light pulse was given and the same time window on the day before, which was used as control. To analyze changes in locomotor activity in response to light, the activity at ZT22 under light (L) was compared to the activity at ZT22 in darkness (D) the day before (third column Fig.4, S3).

Data was analysed with Excel (Microsoft) and Prism 8.2 (GraphPad). The Shapiro-Wilk test was used to analyse normal distribution. Group means were compared by an ordinary one-way ANOVA with Tukey correction and for not normally distributed data by a Kruskal-Wallis test. Prism was used to plot data. Significance levels refer to the raw p values obtained in the statistical tests.

### Immunohistochemistry

Whole heads have been fixed for 2 hours in 4% PFA. After washing in PBS (Phosphate Buffered Saline) specimens have been embedded in 7% agarose and cut into 70-100 μm sections using a vibratome (Leica VT1000S; ^81^). Sections have been washed three times 10 min in 0.3% PBT (PBS with 0.3% TritonX-100), blocked for 1,5 h in 5% normal goat serum in PBT. Afterwards the first antibody solution was incubated overnight at 4°C. Specimens were washed six times and the second antibody solution was added and incubated overnight at 4°C. After another washing step, specimens were mounted in Vectashield (Vector Laboratories) and stored at 4°C until scanning. Probes have been imaged using a LSM 800 (Zeiss). Afterwards the images have been edited for brightness and contrast using FIJI and Adobe Photoshop (V23.5.3).

The following antibodies were used: rabbit-α-GFP (1:1000; life technologies, A11122), 3C11 α-Synapsin (1:50, ^82^), goat-α-rabbit AlexaFluor 488 (1:250, life technologies, A11034) and goat-α-mouse STAR RED (1:250, Abberior, STRED-1001).

## Data Availability

The datasets generated and analysed during the current study are available from the corresponding author on reasonable request.

## Acknowledgments

We thank Claude Desplan, Craig Montell, Christopher Schnaitmann and Ralf Stanewsky for providing flies and Simon Sprecher for sharing antibodies. This work was supported by grants from the German Research Foundation (DFG) to MS (PA3241/2-1) and DP (PA1979/2-1). Stocks obtained from the Bloomington Drosophila Stock Center (NIH P40OD018537) were used in this study.

## Author contributions

MS and DP conceived and designed the experiments. DB, NDF, CM performed the experiments. TT wrote code for analysis. MS wrote the main manuscript text with significant input from DP and CHF. MS and DB prepared the figures. All authors reviewed the manuscript.

## Additional information

The authors declare no competing interests.

