## Supplementary material for "Immediate effects of light on circadian eclosion and locomotor activity depend on distinct sensory input pathways": Fig.S1-S4

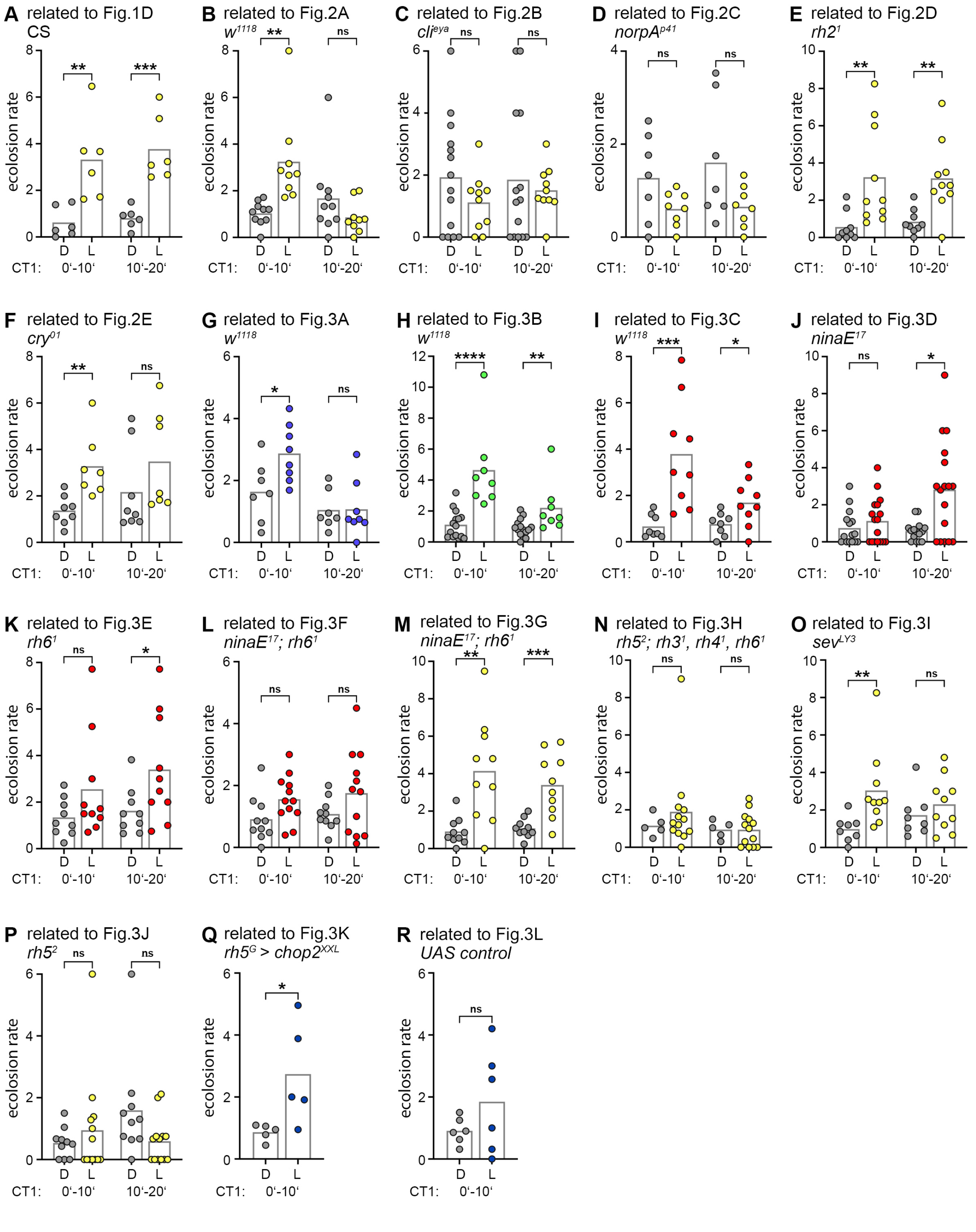


**Fig.S1: Statistical analysis of the immediate light effects on eclosion behaviour. (A-R)** Comparison of the eclosion rate of flies that received a 20 minute light pulse (L) at circadian time (CT) 1 and the appropriate controls kept in darkness (D) in the first 10 min (0'-10') and second 10 min (10'-20') interval. **(A)** Data of Figure 1D. **(B-F)** Eclosion rate of flies shown in Figure 2. **(G-R)** Statistical analysis of data shown in Figure 3. Asterisks denote level of significance: *p≤0.05, **p≤0.01, ***p≤0.001, ****p≤0.0001.

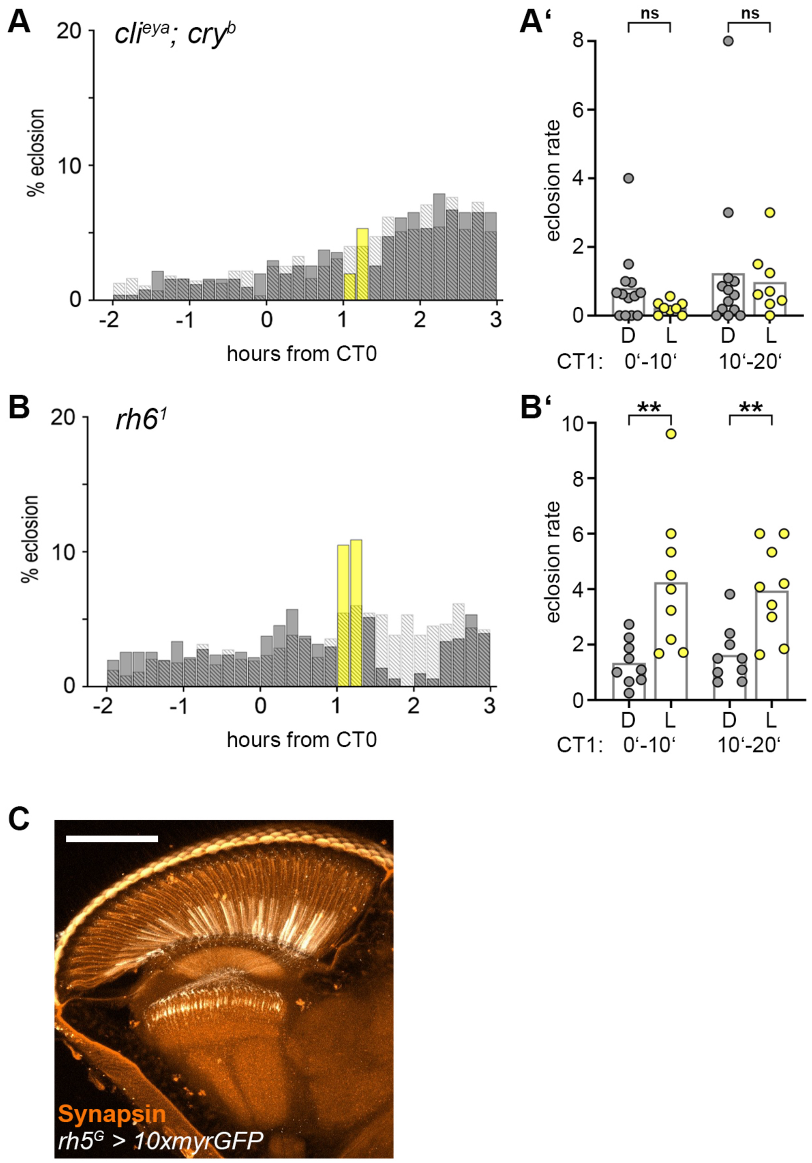


**Fig.S2: The immediate light effect on eclosion behaviour. (A-A')** The light response is gone in flies lacking the compound eyes and CRY function (*cli*$^{eya}$; *cry*$^{b}$). **(B-B')** Flies without Rh6 (*rh6*$^{1}$) respond to light. **(C)** Projection of a head section visualizing Gal4-expression in *rh5-Gal4*-positive cells. The expression can be seen in a subset of R8 cells. Orange = anti-Synapsin; white = anti-GFP, scale bar: 100μm. $n_{exp}$, $n_{ctrl}$ = 506, 549 (A), 504, 731(B). Asterisks denote level of significance: **p≤0.01.

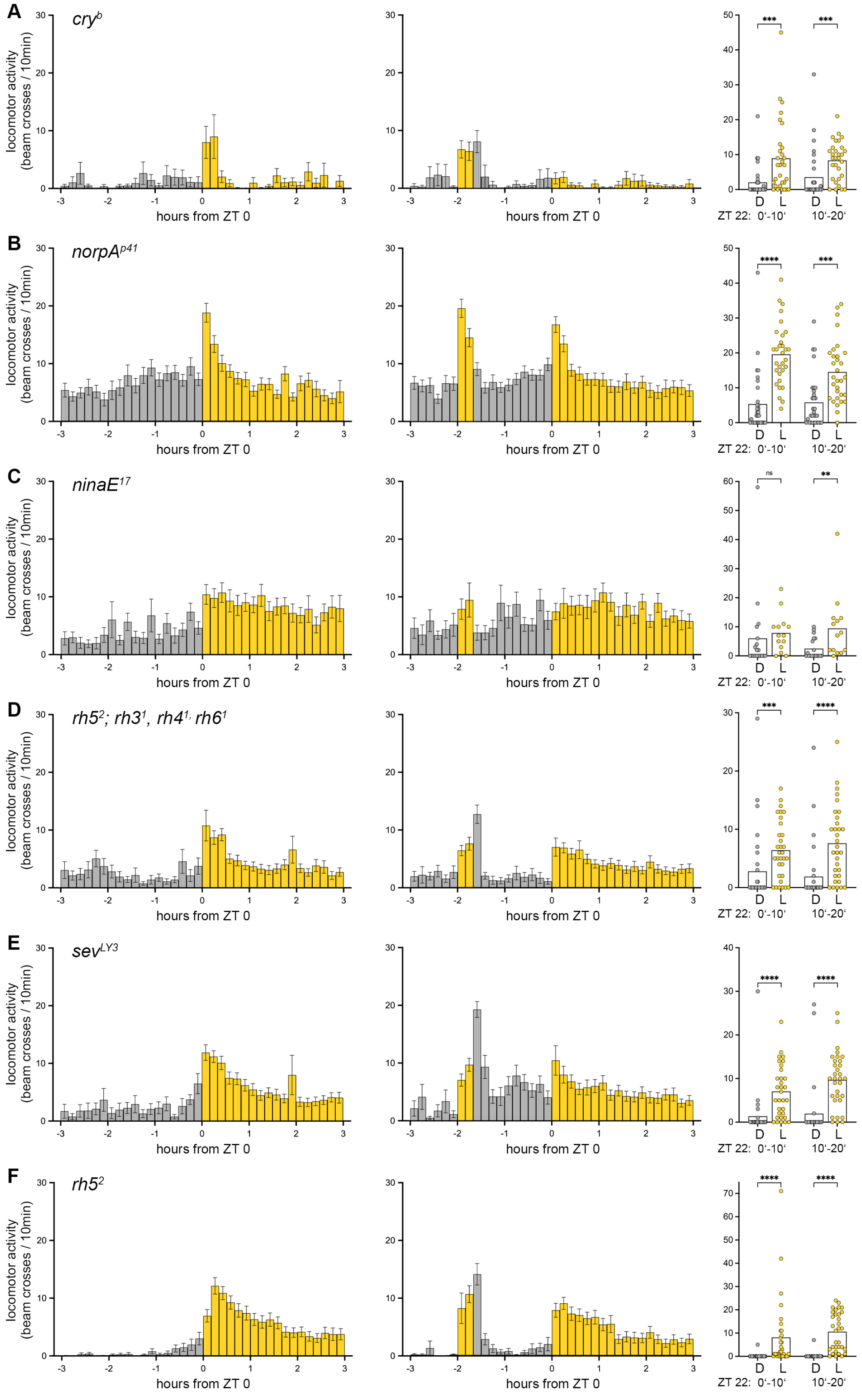


**Fig.S3: The immediate light effect on locomotor activity is visible in flies lacking photosensation in cry-positive or photoreceptor cells. (A-F)** Activity pattern in ten minutes intervals at the time around Zeitgeber time (ZT) 0. First and second column show bar plots of mean ± SEM activity at the day the light pulse was applied (second column) and the activity of the same flies on the previous day (first column). The third column visualizes the comparison between the mean activity at ZT22 in 10 minutes intervals (0'-10' and 10'-20') during the light pulse (L) and the previous control day in darkness (D). All tested flies respond to the light pulse by a significant increase in locomotor activity compared to the appropriate controls. **(A)** Activity data of flies lacking Cryptochrome (cryb), **(B)** flies with impaired phospholipase C activity (norpAp41), **(C)** flies lacking Rh1(ninaE17), **(D)** Rh3, Rh4, Rh5 and Rh6 (rh52; rh31, rh41, rh61), **(E)** R7 (sevLY3) and **(F)** Rh5 (rh52). n = 14- 32; asterisks denote level of significance: $^{**}p \leq 0.01$, $^{***}p \leq 0.001$, $^{****}p \leq 0.0001$.

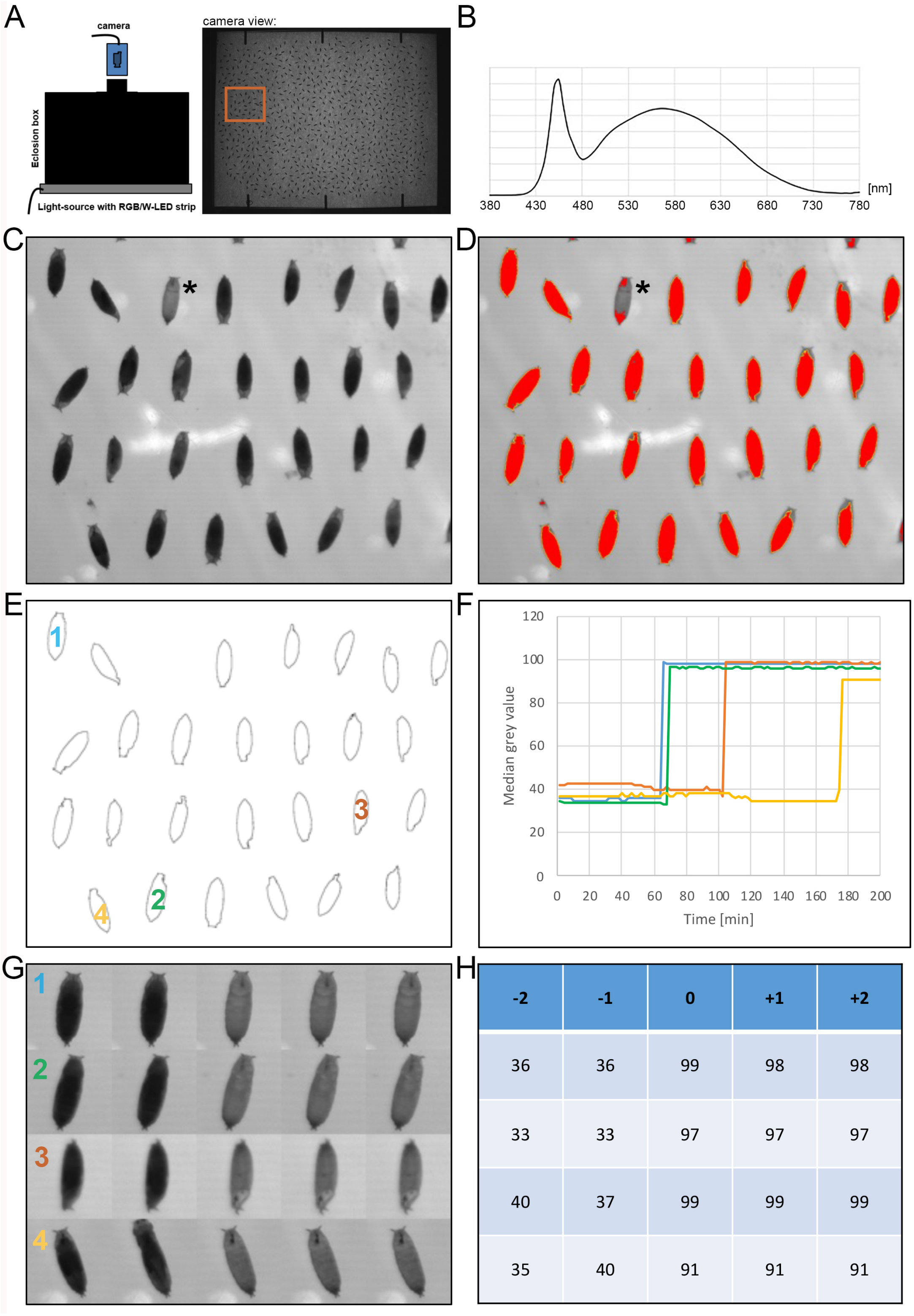


| -2 | -1 | 0 | +1 | +2 |
|---|---|---|---|---|
| 36 | 36 | 99 | 98 | 98 |
| 33 | 33 | 97 | 97 | 97 |
| 40 | 37 | 99 | 99 | 99 |
| 35 | 40 | 91 | 91 | 91 |

**Fig.S4: Analysis of the eclosion monitor data. (A)** Schematic overview of the eclosion box. **(B)** Composition of the white light. **(C)** Several pupae before hatching. Notice the empty pupal case marked with an asterisk. **(D)** Pupae are darker than the background. Pixels with a value below the threshold are coloured red. Yellow outlines mark the different objects (i.e. pupae) that fall within the given size constraints. Notice that the empty pupal chase (marked with an asterisk) is excluded. **(E)** Outlines of the different pupae. Median grey values are calculated of each of these areas over time (i.e. for every frame). The pupae marked 1-4 are shown as examples in D-F. **(F)** Median grey values for pupae 1 to 4 over time. Notice the big jump in brightness from around 40 to around 100 at different points in time. **(G)** Montage of each 2 frames before and after hatching for 4 different pupae. In row 4 the eclosion process can be seen. **(H)** Median grey values for the pupae shown in E.
